# Extracellular Superoxide Dismutase (EC-SOD) Regulates Gene Methylation and Cardiac Fibrosis During Chronic Hypoxic Stress

**DOI:** 10.1101/2020.06.16.154302

**Authors:** Ayan Rajgarhia, Kameshwar Ayyasola, Nahla Zaghloul, Jorge M. Lopez Da Re, Edmund J. Miller, Mohamed Ahmed

## Abstract

**Background:** Chronic hypoxic stress induces epigenetic modifications in cardiac fibroblasts, such as inactivation of tumor suppressor genes (RASSF1A), and activation of kinases (ERK1/2). The effects of the antioxidant enzyme, extracellular superoxide dismutase (EC-SOD), on these epigenetic changes has not been fully explored.

**Objectives:** To define the effect of EC-SOD overexpression on cardiac fibrosis induced by chronic hypoxia.

**Methods:** Wild type C57B6 male mice (WT) and transgenic males with an extra copy of human hEC-SOD (TG) were housed in hypoxia (10% O_2_) for 21 days. Right ventricular tissue was studied for cardiac fibrosis markers using RT-PCR and Western Blot analyses. Downstream effects were studied, for both RASSF-1 expression and methylation and its relation to ERK1/2, using *in-vivo* & *in-vitro* models

**Results:** There were significant increases in markers of cardiac fibrosis : Collagen 1, Alpha Smooth Muscle Actin (ASMA) and SNAIL, in the WT hypoxic animals as compared to the TG hypoxic group (p< 0.05). Expression of DNA methylation enzymes (DNMT 1,2) was significantly increased in the WT hypoxic mice as compared to the hypoxic TG mice (p<0.001). RASSF1A expression was significantly lower and ERK1/2 was significantly higher in hypoxia WT compared to the hypoxic TG group (p<0.05). Use of SiRNA to block RASSF1A gene expression in murine cardiac fibroblast tissue culture led to increased fibroblast proliferation (p<0.05). Methylation of RASSF1A promoter region showed a significant reduction in the TG hypoxic group compared to the WT hypoxic group (0.59 vs 0.75 respectively).

**Conclusions:** EC-SOD significantly attenuates RASSF1A gene methylation, and plays a pivotal role cardiac fibrosis induced by hypoxia.

## Introduction

Fibrosis, in cardiac tissues, can develop following a variety of stimuli, including ischemia, volume overload, pressure overload and hypoxia.^1^ A common feature of all these stimuli is the reduced availability of oxygen. Whether from decreased oxygen delivery or increased oxygen consumption, the tissue hypoxia is associated with infiltration of inflammatory cells and activation of resident cells.^2^ Cardiac fibrosis leads to both systolic and diastolic dysfunction, and increases cell death and damage by inflammatory mediators.^3^

DNA methylation is an epigenetic modification involving alterations in the chromatin structure leading to repression of gene expression. The methylation process is regulated by a family of DNA methyltransferase (DNMT) enzymes. Studies have demonstrated significant increases in the activities of DNMT1 and DNMT 3B in response to hypoxia and the increased activity in these enzymes correlates positively with the degree of cardiac fibrosis. This suggests the role for epigenetic modification in fibrosis secondary to hypoxia.^2^

Hypoxia induced expression of DNMT1 and DNMT 3B is in part regulated by the hypoxia inducible transcription factor 1α, (HIF-1α), through specific hypoxic response elements present in the promoter sequence of DNMT1 and 3B.^2^ Ras association domain family 1 isoform A (RASSF1A) is a tumor suppressor gene and alterations in its regulation are frequently seen in cardiac fibrosis.^2^

RASSF1A functions through its effect on downstream proteins such as extracellular signal regulated kinases (ERK1/2). The Ras/ERK signaling pathway has long been recognized as an intracellular signal transduction critically involved in fibroblast proliferation. DNA methylation mediated silencing of RASSF1A, and subsequent activation of ERK1/2, can lead to activation of fibroblasts and fibroblast proliferation.^4^ Another contributing mechanism to cardiac dysfunction induced by hypoxia is myofilament modification. Two myosin heavy chain (MHC) isoforms, MHC-α and MHC-β are expressed in the mouse heart. α-MHC has higher intrinsic adenosine triphosphatase (ATPase) activity and hence contributes to higher contractility, while β-MHC has lower intrinsic ATPase activity and has a greater economy of force maintenance.^5^ Hypoxia has been shown to cause switching of Myosin Heavy chains (MHC) from its alpha to beta isoform, thereby decreasing ATPase activity and overall force of contraction.^6^

Hypoxia and reactive oxygen species (ROS) play a pivotal role both in the pathogenesis of hypoxia-induced pulmonary hypertension and in the development of cardiac fibrosis.^7–11^ The role of ROS, as a trigger of DNA methylation of tumor suppressor gene promoters in carcinogenesis, was shown in previous studies. The inflammatory response mediated by extracellular reactive oxygen generated from repetitive Ischemia/Reperfusion in a murine model, plays a critical role in the pathogenesis of fibrotic remodeling and ventricular dysfunction.^7^ Supplementation with vitamins E, C, and A have provided antioxidant protection against cardiomyocyte death and have improved survival in congestive heart failure models and doxorubicin-induced injury. *In vitro* cardiomyocyte studies suggest that superoxide dismutase-like, and glutathione peroxidase-like compounds, can protect against free radical production and cellular apoptosis due to doxorubicin.^4^ EC-SOD is an important antioxidant throughout the cardiovascular system. and has been shown to protect the heart from ischemic damage, hypertrophy, and inflammation.^3, 12–13^ Studies show that human populations with a mutation in the matrix-binding domain of EC-SOD, that diminishes its affinity for the extracellular matrix, have a higher risk for the development of cardiovascular and ischemic heart disease.^6^ EC-SOD is important in the prevention of oxidative injury that may contribute to cardiac remodeling, in myocardial infarction models, and alters *ex vivo* heart function.^12–13^ EC-SOD overexpression has also been shown to decrease the fibrosis that develops in cardiac tissue secondary to ischemiareperfusion injury.^8,13^ However, the specific mechanisms by which EC-SOD protects against fibrosis and tissue damage, in various organs including the lung and heart, remain unclear. Also, the relation between EC-SOD and epigenetic changes has not been fully explored. In this study, we reveal the effects of over-expression of EC-SOD on cardiac fibrosis, epigenetic changes and myofilament changes due to chronic hypoxic stress.

## Materials and Methods

### In-vivo studies

All experiments involving animals were reviewed and approved by the Institutional Animal Care and use Committee of the Feinstein Institute for Medical Research. Adult (8 - 10 week old) C57BL6 male mice (wild type - WT) and transgenic neonate mice, with an extra copy of the human EC-SOD gene containing a β-actin promoter (TG), were housed in a pathogen-free environment under standard light and dark cycles, with free access to food and water.^14^ hEC-SOD TG mice with C57BL6 background were studied before, by us and other researchers, and have been well characterized.^14–15^ TG mice act and behave similarly under normal condition (room air), like WT mice strains as shown in many studies before.^14–15^

### Animals

Studied animals were divided into three groups, Group A: WT mice housed in room air; Group B: WT mice housed in 10% normobaric oxygen for 21 days using a BioSpherix chamber (Lacona, NY, USA) (WT hypoxic group). and Group C: TG mice housed under the same hypoxic conditions as Group B.^16^ After 21 days, the animals were euthanized under a surgical plane of anesthesia (Fentany/Xylazine (5:1),^16^ and right ventricular tissue was harvested for analysis.

#### RT-qPCR

Gene expression, within the right ventricular tissue samples, was assessed following tissue disruption and homogenization. RNA was then extracted from the tissue using the AllPrep DNA/RNA extraction kit (Qiagen), according to the manufacturer’s instructions. First strand cDNA synthesis was carried out using SuperScript II RT (Invitrogen). Quantitative real-time PCR primers were designed so that one of each primer pair was exon/exon boundary spanning to ensure that only mature mRNA was amplified. The sequences of the gene-specific primers used were: ASMA, 5’-aatgagcgtttccgttgc 3’ (forward), 5’atccccgcagactccatac 3’(reverse); collagen 1a 1 (COL1A1), 5’ catgttcagctttgtggacct 3’ (forward),5’ gcagctgacttcagggatgt 3’ (reverse); collagen 3 a 1 (COL 3A1), 5’ tcccctggaatctgtgaatc 3’ (forward), 5’tgagtcgaattggggagaat 3’ (reverse); α-Myosin Heavy Chain (Myh6) 5’ cgcatcaaggagctcacc-3’ (forward), 5’-cctgcagccgcattaagt-3’ (reverse), β-Myosin Heavy Chain (Myh7) 5’-cgcatcaaggagctcacc - 3’ (forward), 5’ctgcagccgcagtaggtt-3’ (reverse). Q-PCR was performed; amplification and detection were carried out using Roche Applied Science LightCycler480 PCRSystems with software. The PCR cycling program consisted of 45 three-stepcycles of 15 s/95 °C, 1 min/57 °C, 1 sec/72 °C. Relative changes in mRNA expression were calculated as fold-changes (normalized using *Gapdh*) by using the comparative Ct (ΔΔCt) method.^17^

#### Immunohistochemistry - Collagen 1 (Cy 3)

Right ventricular tissue, fixed in 4% paraformaldehyde for 24 h, was processed, embedded in paraffin, and subsequently cut into 4μm-thick sections. The slides then underwent heat mediated antigen retrieval followed by incubation with primary antibody - anti-Collagen 1 antibody-3G3- (Abcam - ab88147) and the secondary antibody - AffiniPure Goat Anti-Mouse IgG (Jackson Immunoresearch Lab Inc - Code - 115005003). The slides were then analyzed using an Olympus FluoView FV300 Confocal Laser Scanning Microscope (Thermo Fisher Scientific) and the Fiji image processing software, (an open source platform for biological image analysis), was used for analysis of pixel density.^18^

#### Western Blot analysis

Frozen right ventricular tissues were homogenized, and protein extraction was carried out using a total protein extraction Kit (BioChain Institute, Inc. Hayward, CA). Protein concentration was evaluated using the Modified Lowry Protein Assay (Thermo Fisher Scientific, Rockford, IL, USA). Samples were prepared for SDS-PAGE in Laemmli sample buffer (Bio-Rad, Hercules, CA, USA) and processed as previously described.^18^ Membranes were briefly washed and immediately incubated with the respective primary antibody in 5% BSA with phosphate buffered saline with Tween 20 (PBST), overnight. Following washing with PBST, the membranes were incubated with horseradish peroxidase - conjugated secondary antibodies for 40–60 min. The membranes were washed and then processed using Amersham ECL detection systems (GE healthcare, Piscataway, NJ USA). The membranes were immediately exposed to 8610 Fuji X-Ray Film. The films were assessed using Quantity One 1-D Analysis Software on a GS-800 Calibrated Densitometer. The density of each band is presented as a ratio in comparison to the Actin band density. Primary antibodies were used to detect the following markers: Collagen 1, Alpha Smooth Muscle Actin, SNAIL, DNMT1 and 3b, HIF1α (each obtained from Abcam, (Cambridge, MA); RASSF1A (Origene, Rockville, MD,), and ERK 1/2 (Cell Signaling Technology, Danvers, MA, USA)

#### ROS assay

Intracellular ROS was assessed using a cell-based assay for measuring hydroxyl, peroxyl, or other reactive oxygen species. The assay employs the cell-permeable fluorogenic probe 2’, 7’ Dichloro-dihydrofluorescin diacetate (DCFH-DA) (Cell Biolab., San Diego, CA).

### In-vitro analysis

Primary C57BL6 Mouse Cardiac Fibroblasts (MCF) (Cell Biologics, Chicago, IL, Catalog No. C57-6049) were grown to confluence in T75 flasks using Fibroblast growth medium (Cell Biologics, Chicago, IL, Catalog No. M2267), in a humidified incubator (37°C, 5% CO_2_). Cells were seeded to six-well tissue culture plates, at a density of 10,000 cells per well and incubated for 24 h. Next, the ells were transfected with human EC-SOD (hEC-SOD) cDNA inserted into a vector plasmid pcDNA3 (5446 nucleotides; Invitrogen Life Technologies, Carlsbad, CA, USA) as previously described,^14^ using the FuGENE kit (Roche Diagnostics, Indianapolis, IN, USA). Each well received 1 μg DNA/100μL serum free medium of the DNA/FuGENE complex. Control wells received serum free medium alone. Transfected cells were selected using Geneticin (Invitrogen Life Technologies). Transfection of the fibroblasts was confirmed by Western blot analysis using an antibody specific for hEC-SOD (R&D Systems, Minneapolis, MN, US)

#### Quantitative Flow Cytometry - Effects of hypoxia on Global DNA Methylation profile

A modular incubator chamber (Billups-Rothenberg, Del Mar, CA, USA) was used for the cell hypoxia studies and a 1% oxygen atmosphere maintained using an oxygen sensor (BioSpherix, Lacona, NY, USA). Cells were maintained in a microenvironment of 37°C, 1% O2, 5% CO2 and 100% humidity. To examine the effects of hypoxia ± EC-SOD overexpression on global DNA methylation, both transfected and non-transfected MCF were incubated in hypoxia for 72 h. Control, non-transfected MCF, were maintained in 21% oxygen for 72 h. Post culture, MCF cells were fixed in Carnoy’s solution prior to 60 min acid hydrolysis in 1 M HCl at 37°C. Following this DNA denaturation step, cells were then treated with either an anti-5’ methylcytidine (5MeC) monoclonal antibody (Epigentek, Catalog No. A-1014), or non-specific IgG1 (BD Biosciences, San Jose, CA). IgG1-negative controls were used at the same concentration as the primary antibody. Immunostaining was conducted using an FITC-conjugated rabbit anti-mouse secondary antibody (Thermo Scientific, Catalog No. 31561). The cells were then subjected to flow cytometry (BD Biosciences, San Jose, CA) and the results assessed using CellQuest Pro (BD Biosciences, San Jose, CA).

#### Proliferation Studies - Effects of Blocking RASSF1A expression

Expression of RASSF1A was reduced by transfection of MCF with Small interfering RNA (SiRNA) SiRASSF1A (Thermo Fisher Scientific, catalog no. 185488) located on Chr.9: 107551555 - 107562267 on Build GRCm38MCF, using Lipofectamine RNAiMAX tranfection protocol (Life Technologies). Effectiveness of the transfection was evaluated at 24h, 48h. 72h and 96hr post transfection with western Blot analysis using an antibody specific for RASSF1A (Abcam). Posttransfection studies showed that RASSF1A expression is minimal after 72hr of transfection (data are not shown). This time point was used for cell proliferation assessment in the next step. Cells were housed in a humidified incubator (37°C, 5% CO_2_) and compared to control MCF, which were transfected with an empty vector and kept under the same conditions. After 72hr of transfection, MCF proliferation was assessed by BrdU (5-Bromouridine) incorporation, (Roche Diagnositic, Mannheim) Cell counts were performed using a hemocytometer 24 and 48 hours later.

#### Methylation study

Bisulfite chemically converts unmethylated cytosine to uracil but has no effect on methylated cytosine. To determine if promoter region hyper-methylation could be responsible for the down-regulation of RASSF1A expression, direct bisulfite sequencing on mouse cardiac fibroblasts, was performed.^18^ The methylation ratio (meth-ratio) was calculated using methylated CpG total CpG counts.

### RASSF1A Methylation Analysis

Briefly, the genomic DNA from mouse heart was isolated using the Trizol solution (Invitrogen, Thermo Fisher Scientific Inc), according to the manufacturer’s protocols. The methylation status of the RASSF1A promoter region was determined by chemical modification of genomic DNA with sodium bisulfite and methylation-specific PCR. The bisulfite-treated DNA was used as a template for the methylation-specific PCR reaction. Primers for the unmethylated DNA-specific reaction were: F, 5V-GGTGTTGAAGTTGTGGTTTG-3V; R, 5V-TATTATACCCAAAACAATACAC-3V. Primers for the methylated DNA-specific reaction were: F, 5V-TTTTGCGGTTTCGTTCGTTC-3V; R, 5V-CCCGAAACGTACTACTATAAC-3V. The reactions were incubated at 95°C for 1 minute, 55°C for 1 minute, and 72°C for 1 minute, for 35 cycles. The amplified fragment was confirmed by DNA sequencing. DNA from normal heart was used as a control for unmethylated RASSF1A. The strategies for RASSF1A sequencing and the amplicons map Supplements 1 and 2 respectively.

### Statistics

Data was expressed as the mean ± standard error of the mean (SEM). Unless otherwise indicated, a one-way or two-way analysis of variance followed by Bonferroni post hoc test was used to assess significance (P<0.05) using GraphPadprism 8 (GraphPad Software, Inc, La Jolla, CA).

## Results

A comparison between WT adult mice and TG mice housed in room air was performed, and shown in Suplement (#3). All molecular testing showed no significant difference between the two strains under normoxic conditions.

### EC-SOD reduces cardiac fibrosis

#### Gene expression analysis of markers of fibrosis

Gene expression analysis using RT-PCR showed a significant decrease in the expression of Collagen 1 (Col1A1) (Fig. 1A), Collagen 3 (Fig. 1B) and ASMA (Fig 1C) in the hypoxic TG, as compared to the hypoxic WT animals (p<0.05). Gene expression of Collagen 1, Collagen 3 and ASMA in hypoxic TG group was not statistically significantly different from room air controls. These results show that EC-SOD over-expression reduces gene expression of cardiac fibrosis markers to levels comparable to RA groups.

**Figure 1:**
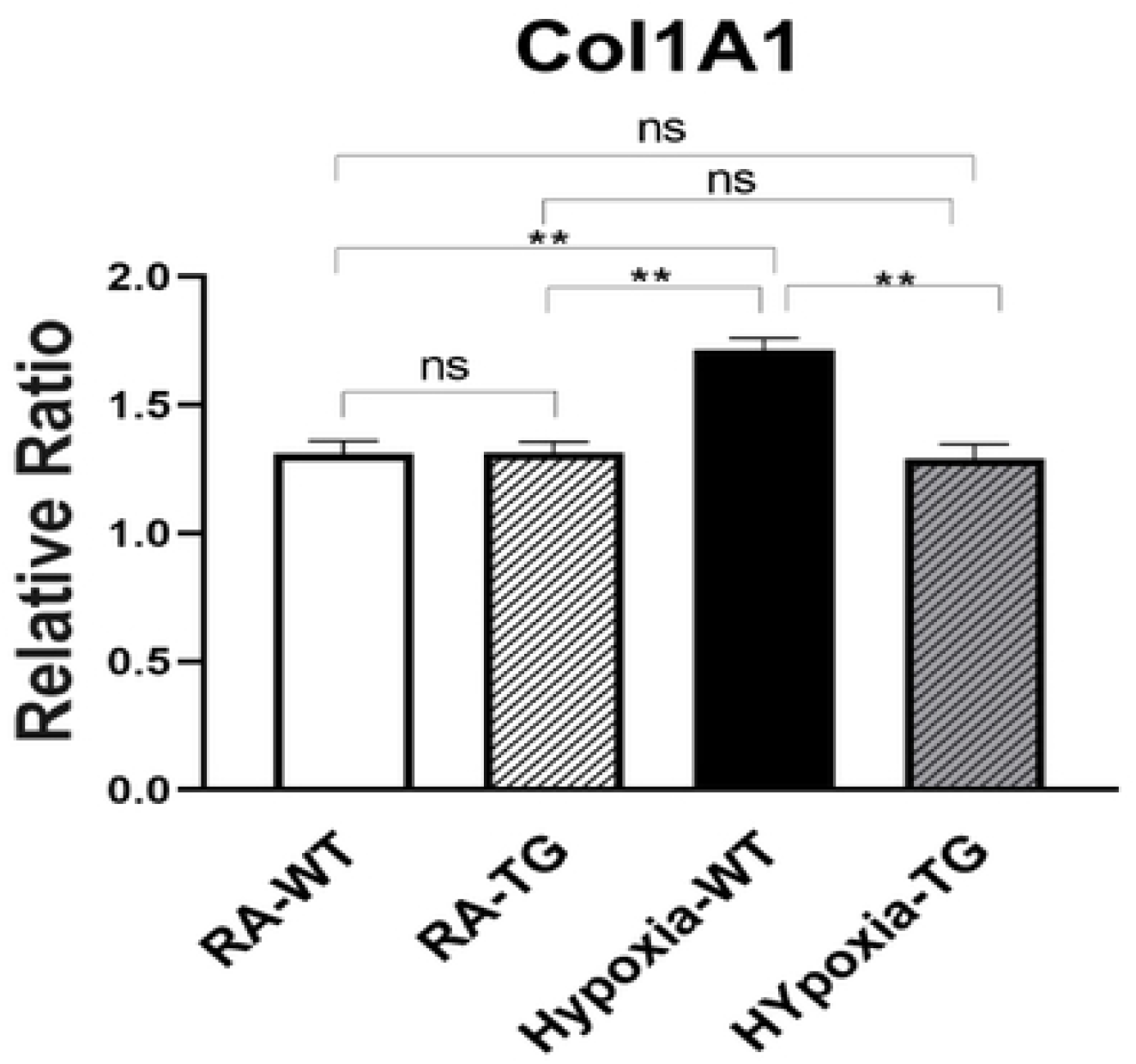
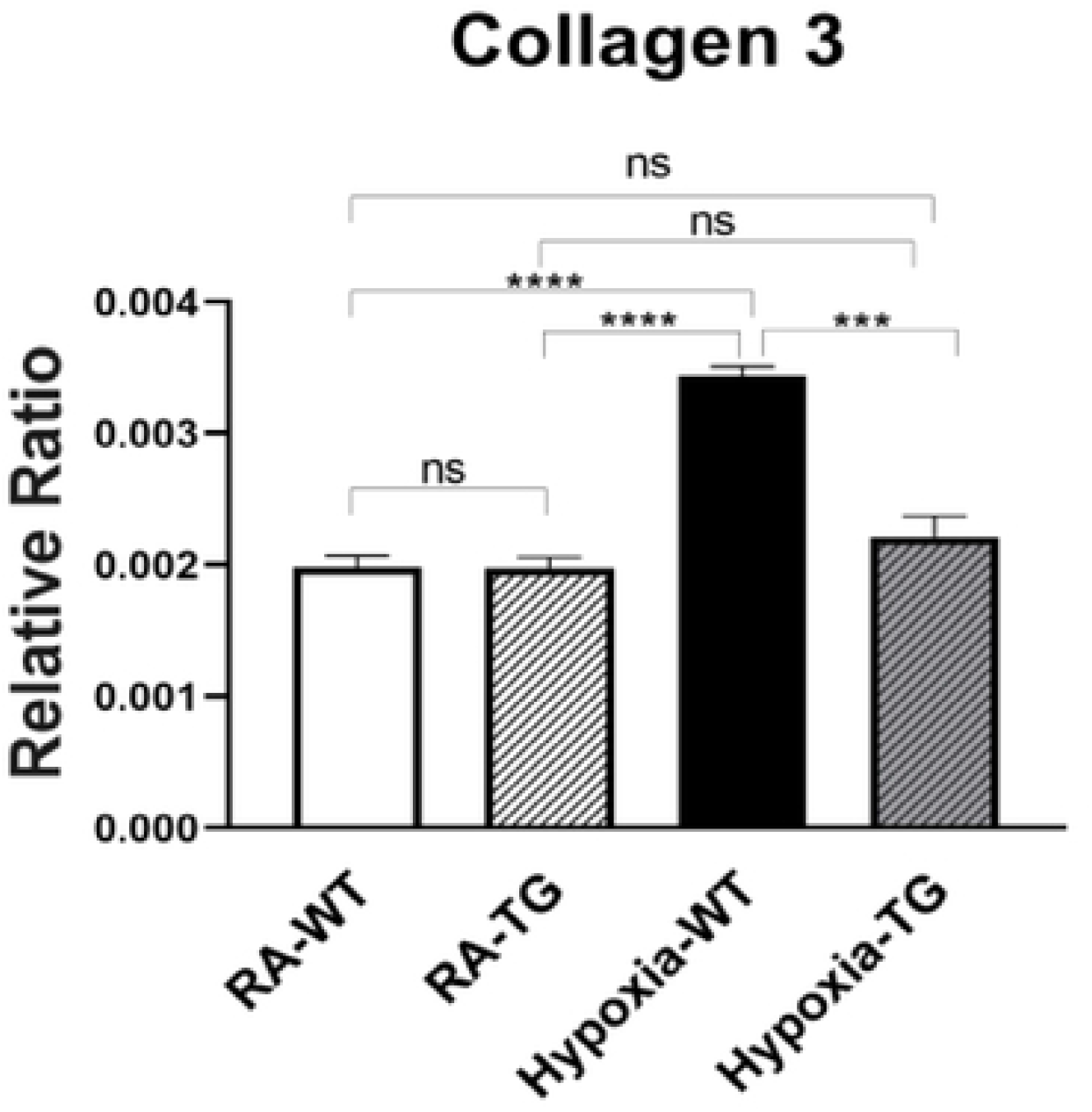
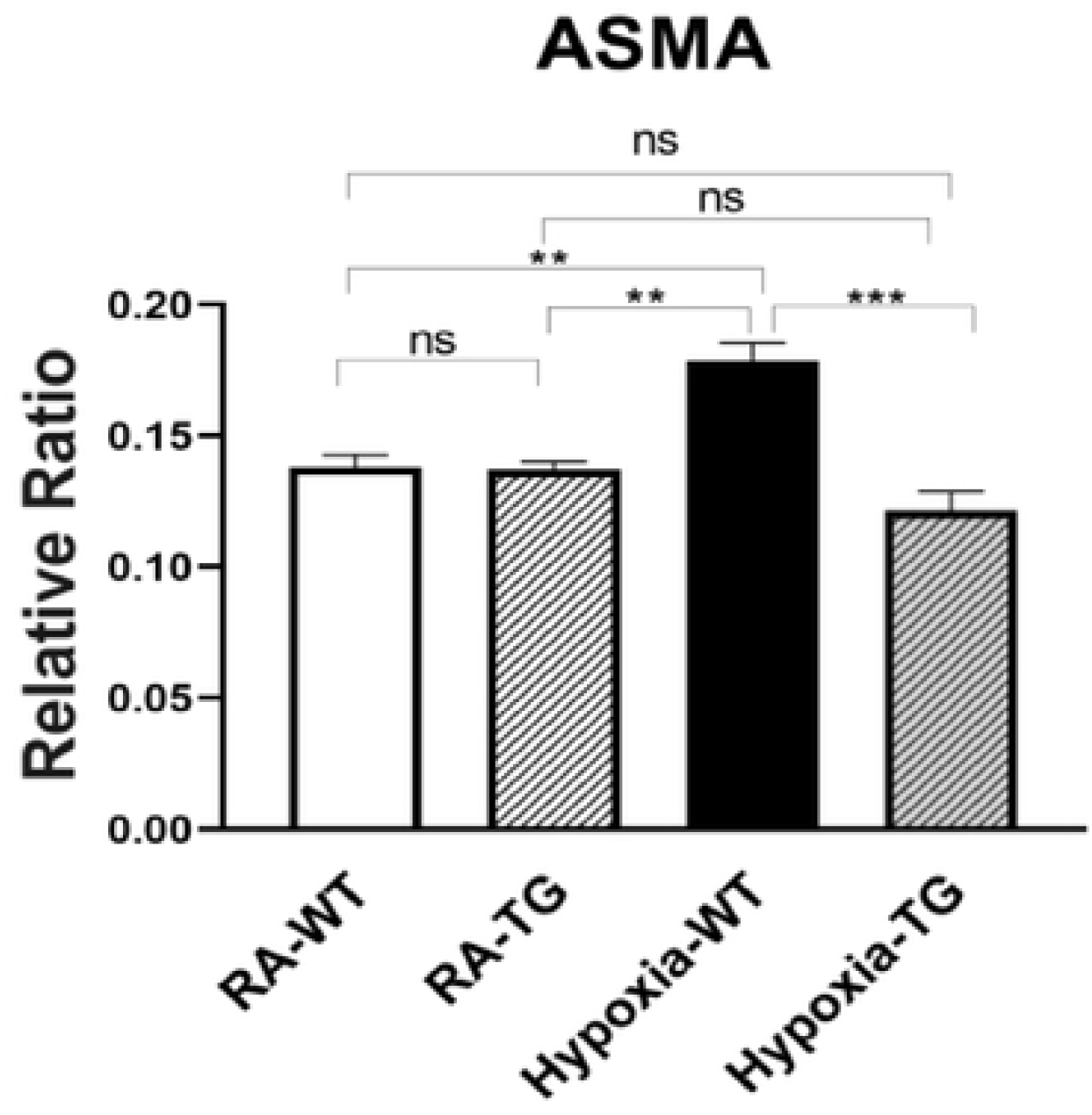
RT-PCR for markers of fibrosis - Collagen 1, Collagen 3 and ASMA. Adult male C57B6 mice, WT and EC-SOD transgenic mice (TG), were exposed to FiO_2_ 10% hypoxia for 21 days (WT). The control group was composed of animals housed in room air (RA). Quantitative RT-PCR was used to assess gene expression analysis of Collagen 1 (A), Collagen 3 (B) and ASMA (C) in the right ventricular tissue. All experiments n = 5. Data represents mean + SEM. *P:0<05, ** P< 01, *** P< 0.001, **** P < 0.0001.

#### Protein levels of markers of fibrosis

Protein level assessment using Western blot analysis showed a significant increase in the levels of Collagen 1 (Fig 2A), ASMA (Fig 2B) and SNAIL1 (Fig 2C) in the hypoxic WT animal as compared to the hypoxic TG animals (p<0.05). Protein concentration of these three markers in the TG animals was not significantly different from the RA control group. This indicates that EC-SOD overexpression reduces protein expression of cardiac fibrosis markers to levels comparable to RA groups.

**Figure 2:**
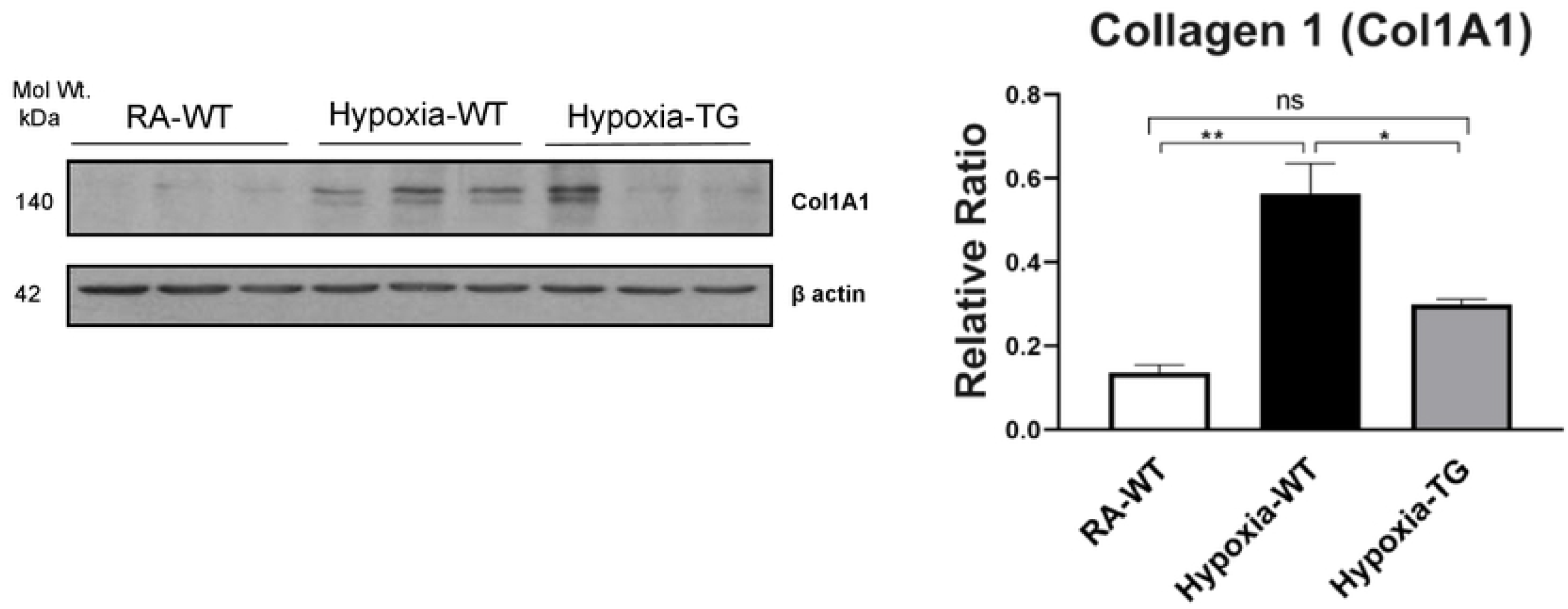
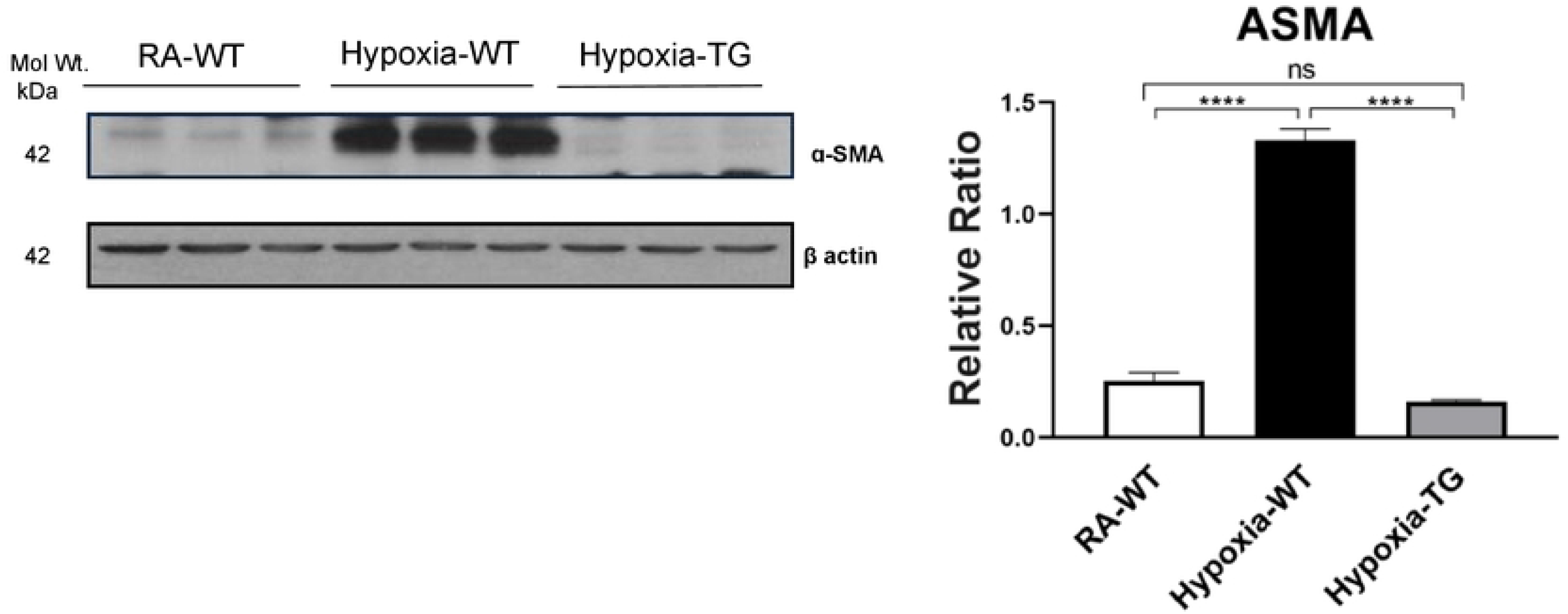
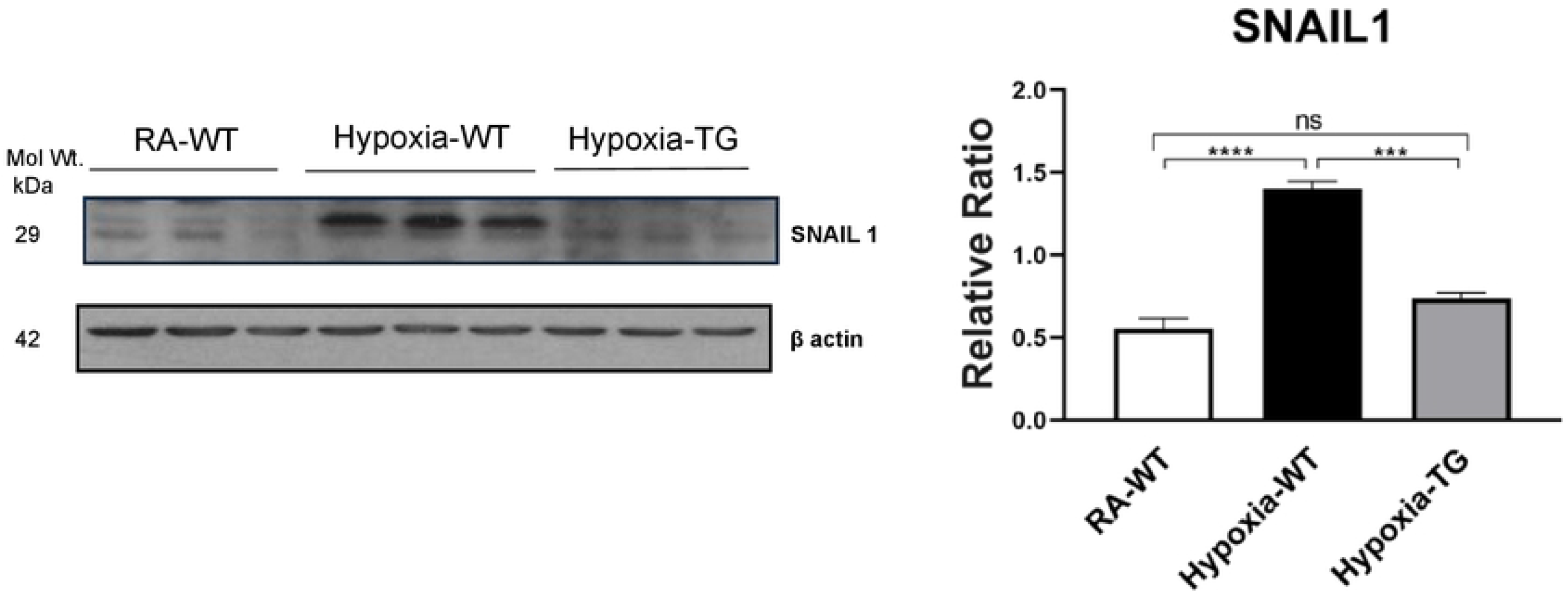
Western Blot Analysis of markers of fibrosis - Collagen 1, ASMA and SNAIL. Adult mice (C57B6) (WT) and EC-SOD transgenic mice (TG), were exposed to FiO2 10% hypoxia for 21 days (WT). Room air animals were used as control group (RA). WB analysis was used to assess protein levels of Collagen 1 (A), ASMA (B) and SNAIL (C) in the right ventricular tissue. All experiments n=3. Data represents mean + SEM. *P:0<05, ** P< 01, *** P< 0.001, **** P < 0.0001.

#### Immunohistochemistry staining for markers of fibrosis

Immunohistochemistry staining for Collagen 1 showed a significant decrease in pixel density of Collagen 1 in the hypoxic TG animals, as compared to the hypoxic WT animals (Fig 3A&B) (P<0.05).

**Figure 3:**
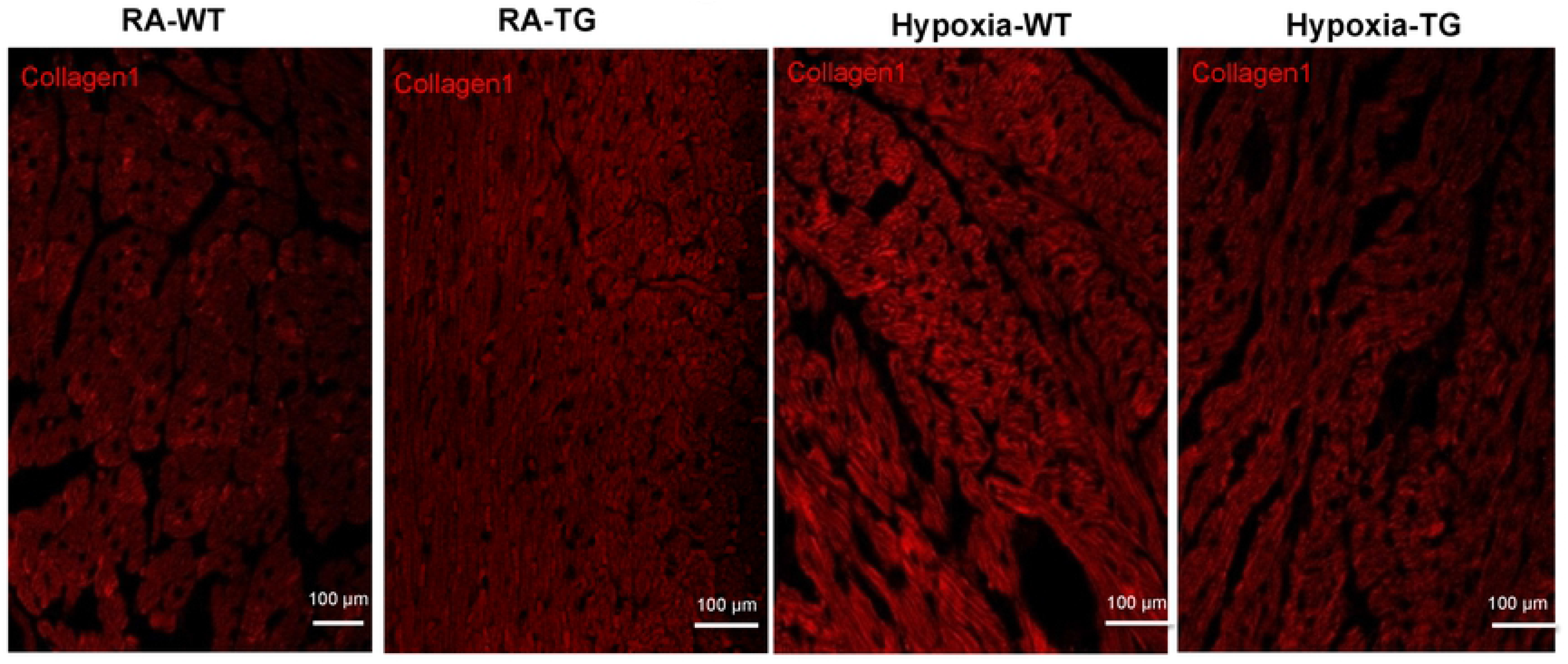
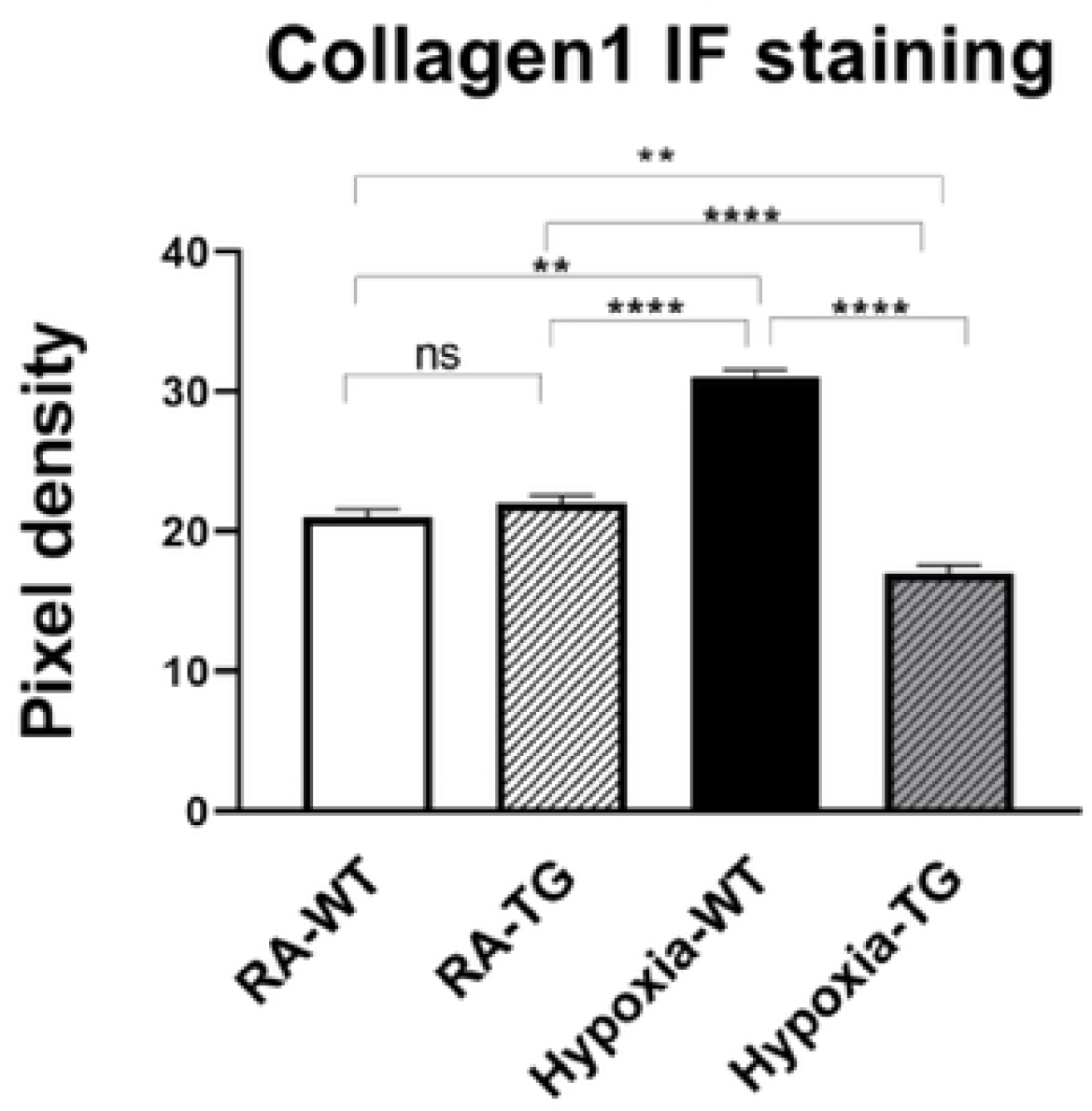
Immunohistochemistry for Collagen 1. Right ventricular tissue from adult wild type mice WT and EC-SOD transgenic mice (TG), which were exposed to FiO_2_ 10% hypoxia for 21 days (WT) and a room air control group (RA), were treated with Cy3 stain to assess for levels of Collagen 1 (Fig 3A). The images were analyzed using Fiji image processing software to measure pixel density (Fig 3B). All experiments n=5. Data represents mean + SEM. *P:0<05, ** P< 01, *** P< 0.001, **** P < 0.0001.

### EC-SOD reduces epigenetic modifications - DNA Methylation

#### Flow Cytometry analysis of global DNA Methylation

Quantitative flow cytometry studies, using antibody directed to methylated DNA, revealed a significant increase in global DNA methylation in cardiac fibroblasts subjected to hypoxia, compared to cardiac fibroblasts transfected with hEC-SOD and subjected to the same hypoxic conditions (p< 0.05) (Fig. 4).

**Figure 4:**
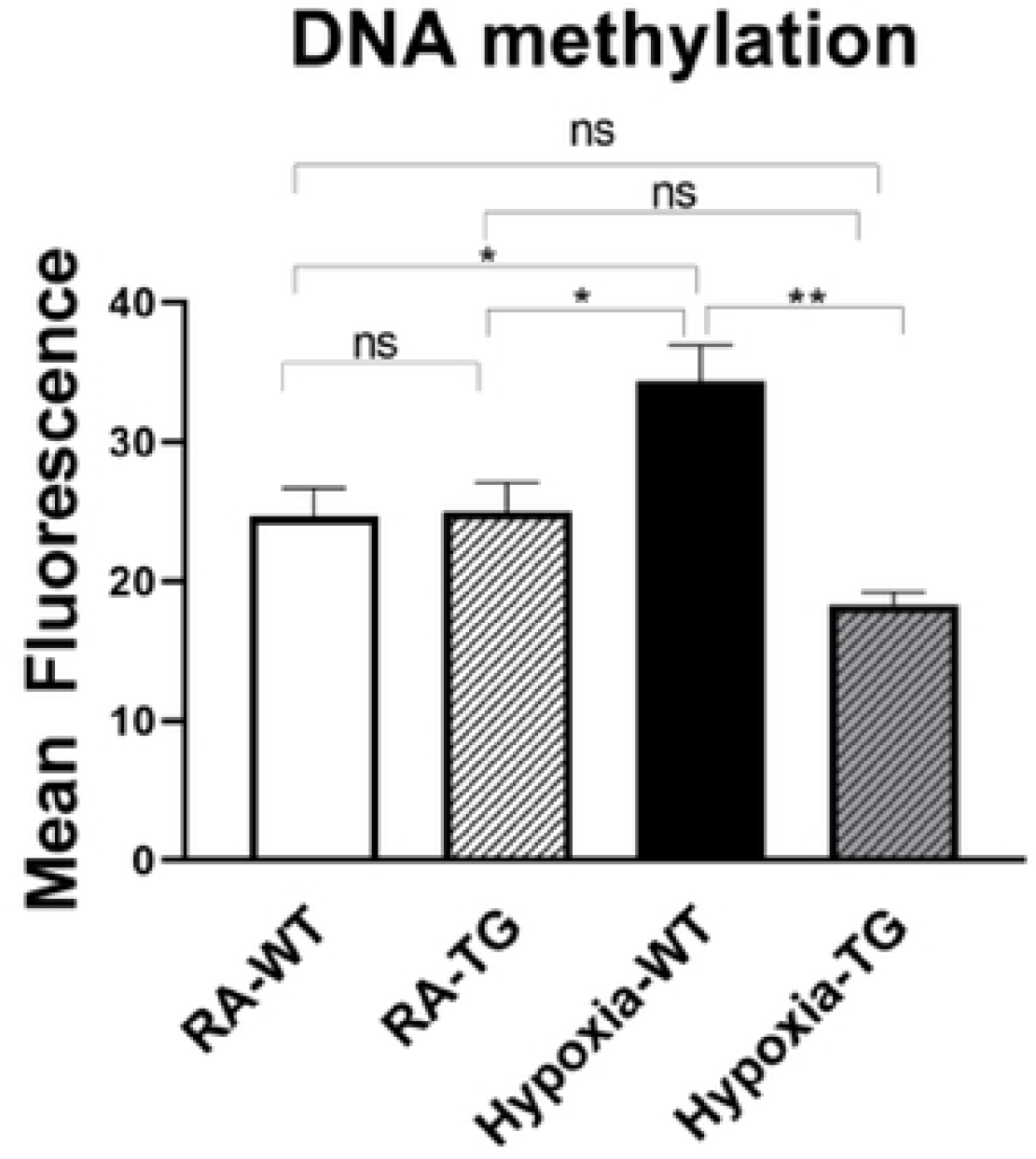
Flow Cytometry studies for DNA Methylation –. **In** in-vitro studiesusing cardiac fibroblasts, DNA methylation levels were assessed in cells exposed to hypoxic conditions (FiO_2_ 1% for 72 hours) and compared to cells transfected with EC-SOD and exposed to hypoxia. Cells cultured in room air were employed as controls. Triplicate wells were analysed, and the experiment were repeated 5 times. Data represents mean + SEM. *P:0<05, ** P< 01, *** P< 0.001, **** P < 0.0001.

#### Protein levels of DNMT 1, 3b and HIF1α

Protein assessment using Western blot analysis showed a significant decrease in DNMT1 (Fig 5A) and DNMT-3b (Fig 5B) in the hypoxic TG animals compared to the hypoxic WT animals (p<0.05). The levels of HIF1α (Fig 5C) were also noted to be significantly reduced in the hypoxic TG animals when compared to the WT animals in hypoxia (p<0.05). However, the levels in the hypoxic TG animals were not significantly different when compared to the TG room air group. Assay of free reactive oxygen species accumulation by DCF assay, showed a significant increase of ROS in the WT hypoxic group, but was significantly lower in the TG hypoxic group (Fig. 5D).

**Figure 5:**
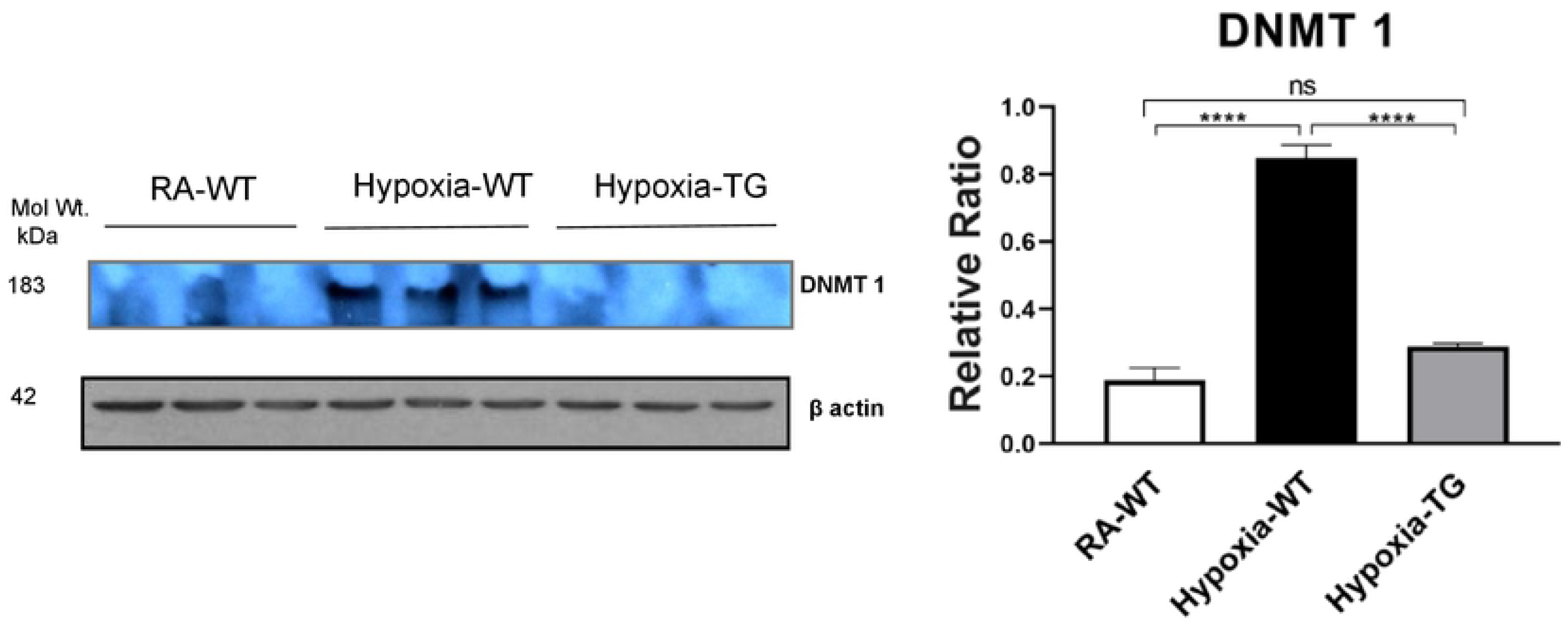
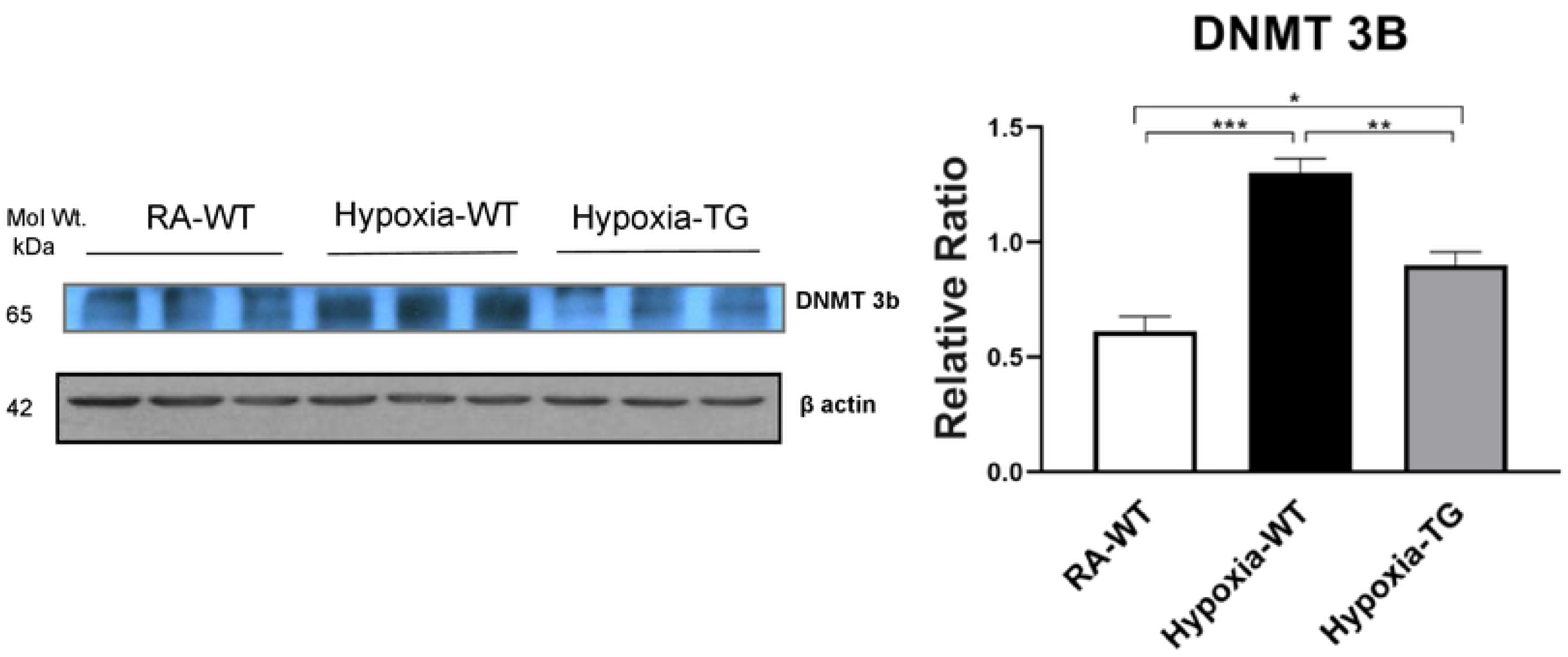
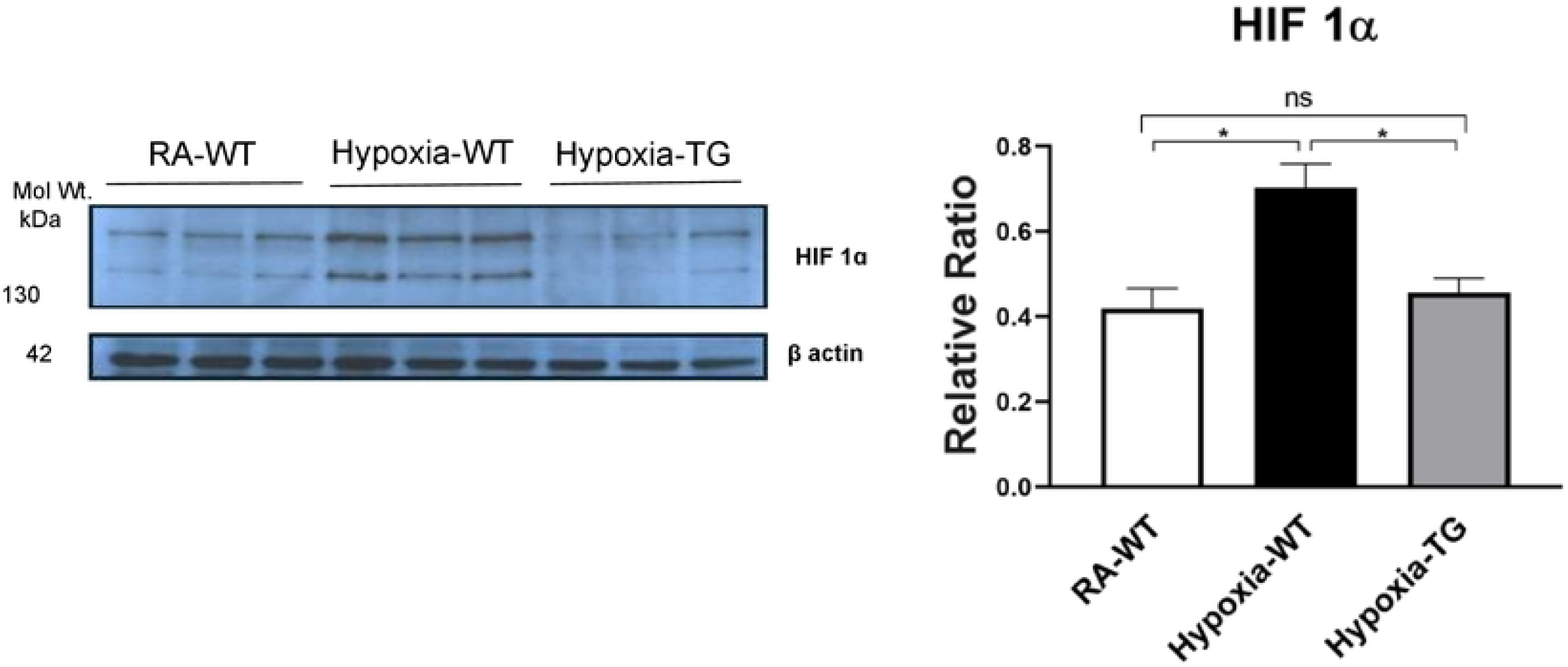
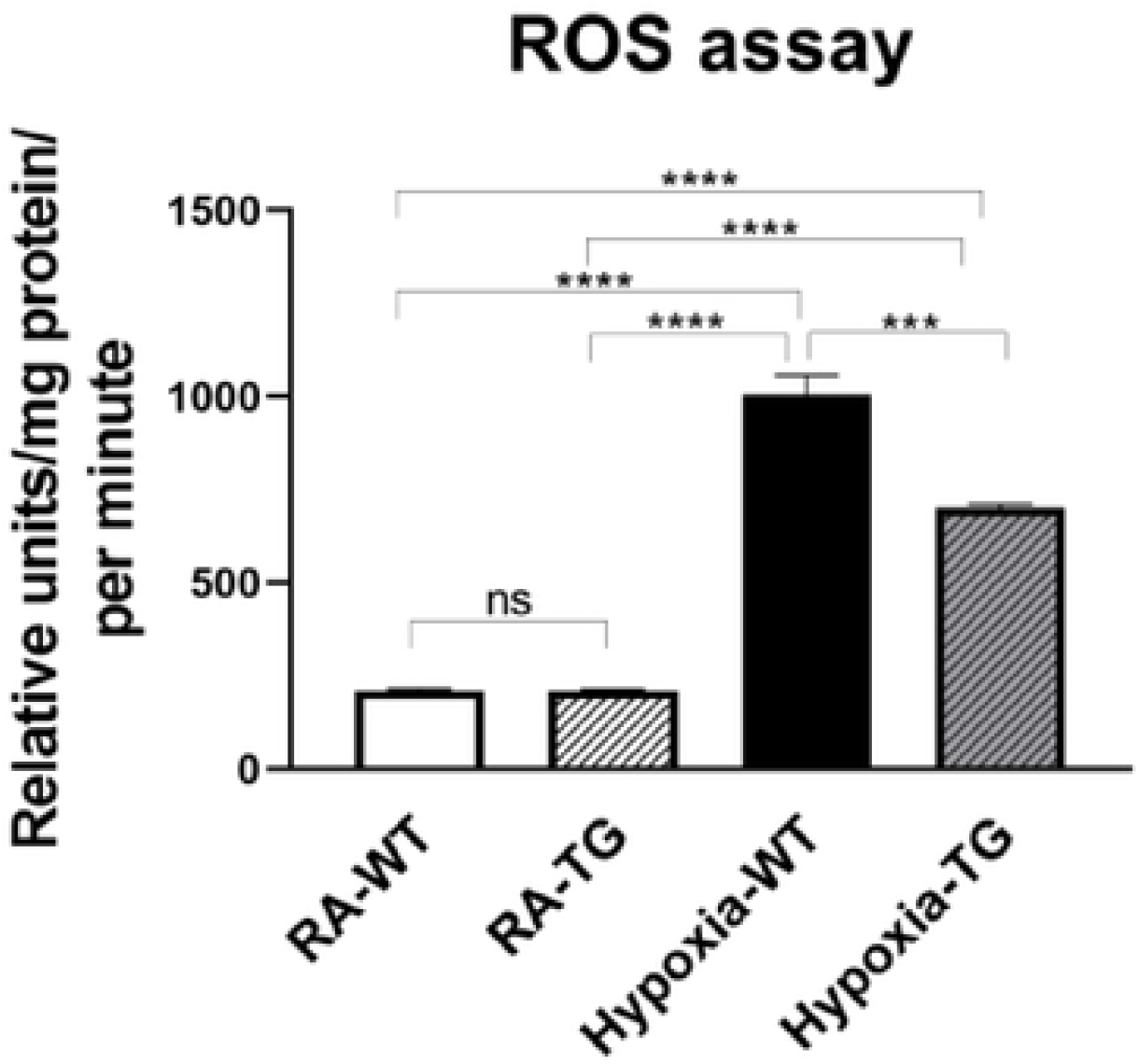
Western Blot analysis for DNMT1, 3b and HIF-1α. Adult mice (WT) and EC-SOD transgenic mice (TG), were exposed to FiO_2_10% hypoxia for 21 days . Room air animals were used as **a** control group (RA). WB analysis was used to assess protein levels of DNMT 1 (A); DNMT 3b (B); HIF-1α assessment in cardiac fibroblasts (C) in the right ventricular tissue. ROS assay in all studied groups (D). All experiments were carried out in triplicate. Data represent mean + SEM. *P:0<05, ** P< 01, *** P< 0.001, **** P < 0.0001.

### Hypoxia induced epigenetic modifications - RASSF1A and ERK 1/2

#### Protein levels of RASSF1A and ERK1/2

Western Blot analysis in the hypoxic WT animals showed a significant reduction in the gene RASSF1A (Fig 6A), as compared to both hypoxic TG animals and RA control groups (p< 0.05). WT hypoxic animals showed a significant increase of ERK phosphorylation in comparison to RA control group (P<0.05). The level of ERK 1/2 (Fig 6B) was significantly reduced in the hypoxic TG animals compared to the hypoxic WT animals.

**Figure 6:**
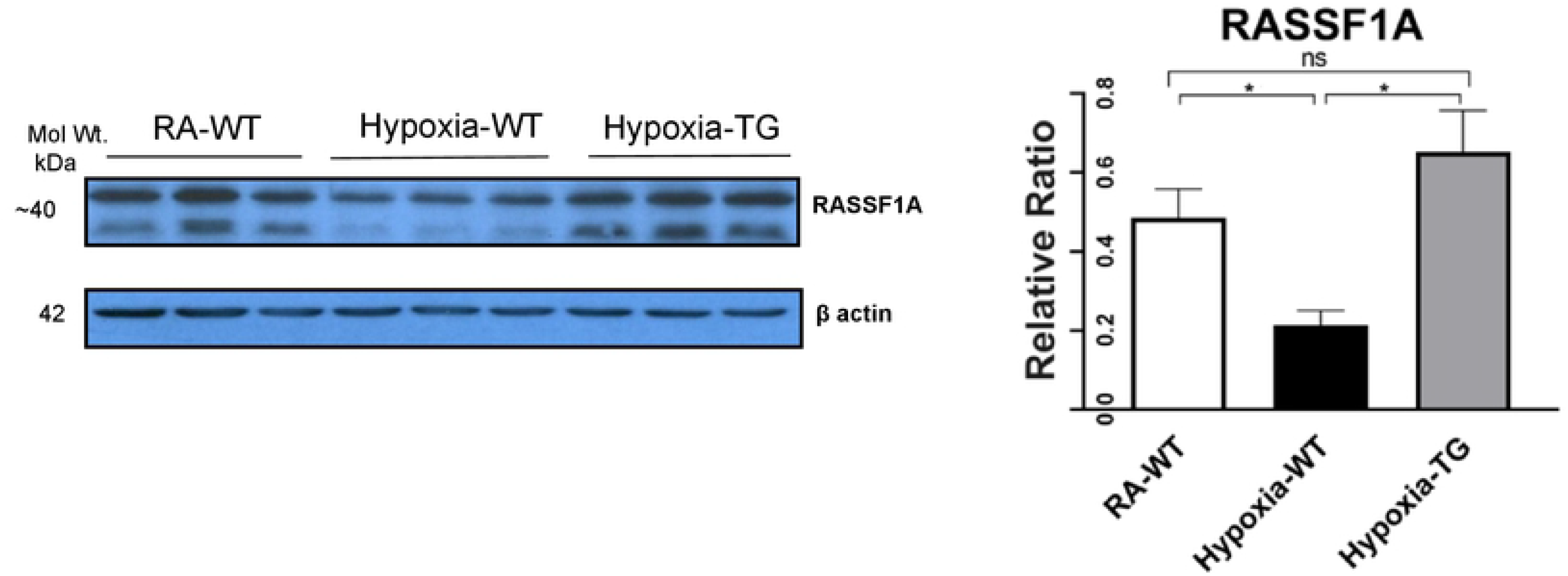
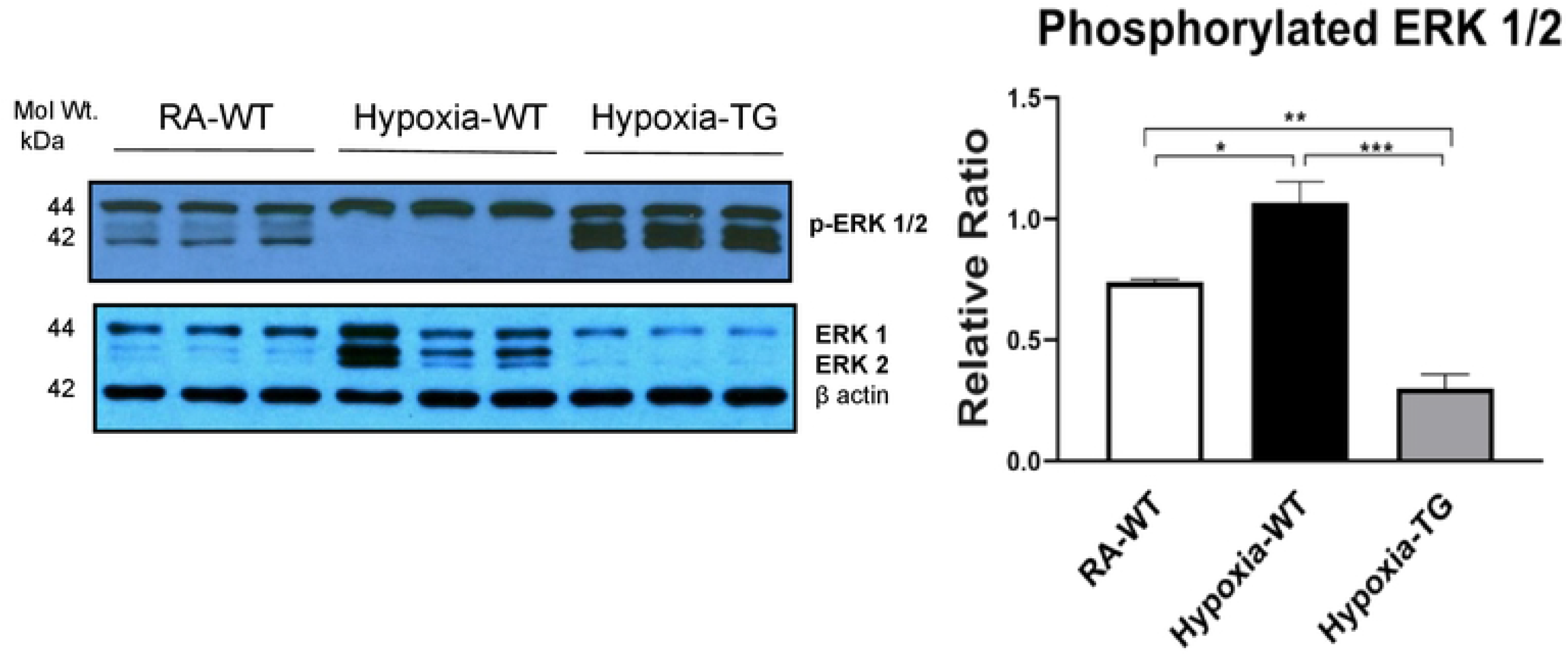
Western Blot analysis for RASSF1A and ERK1/2. Adult mice (C57B6) (WT) and EC-SOD transgenic mice (TG), were exposed to 10% hypoxia for 21 days (WT). Room air animals were used as control group (RA). WB analysis was used to assess protein levels of RASSF1A (A) and ERK1/2 (B) in the right ventricular tissue. All experiments were carried out in triplicate. Data represents mean + SEM. *P:0<05, ** P< 01, *** P< 0.001, **** P < 0.0001.

#### EC-SOD prevents myofilament changes

Gene expression analysis using RT-PCR showed a significant reduction in the levels of α-MHC (Fig. 7) in hypoxic WT animals as compared to RA groups and hypoxic TG group (P<0.05).

**Figure 7:**
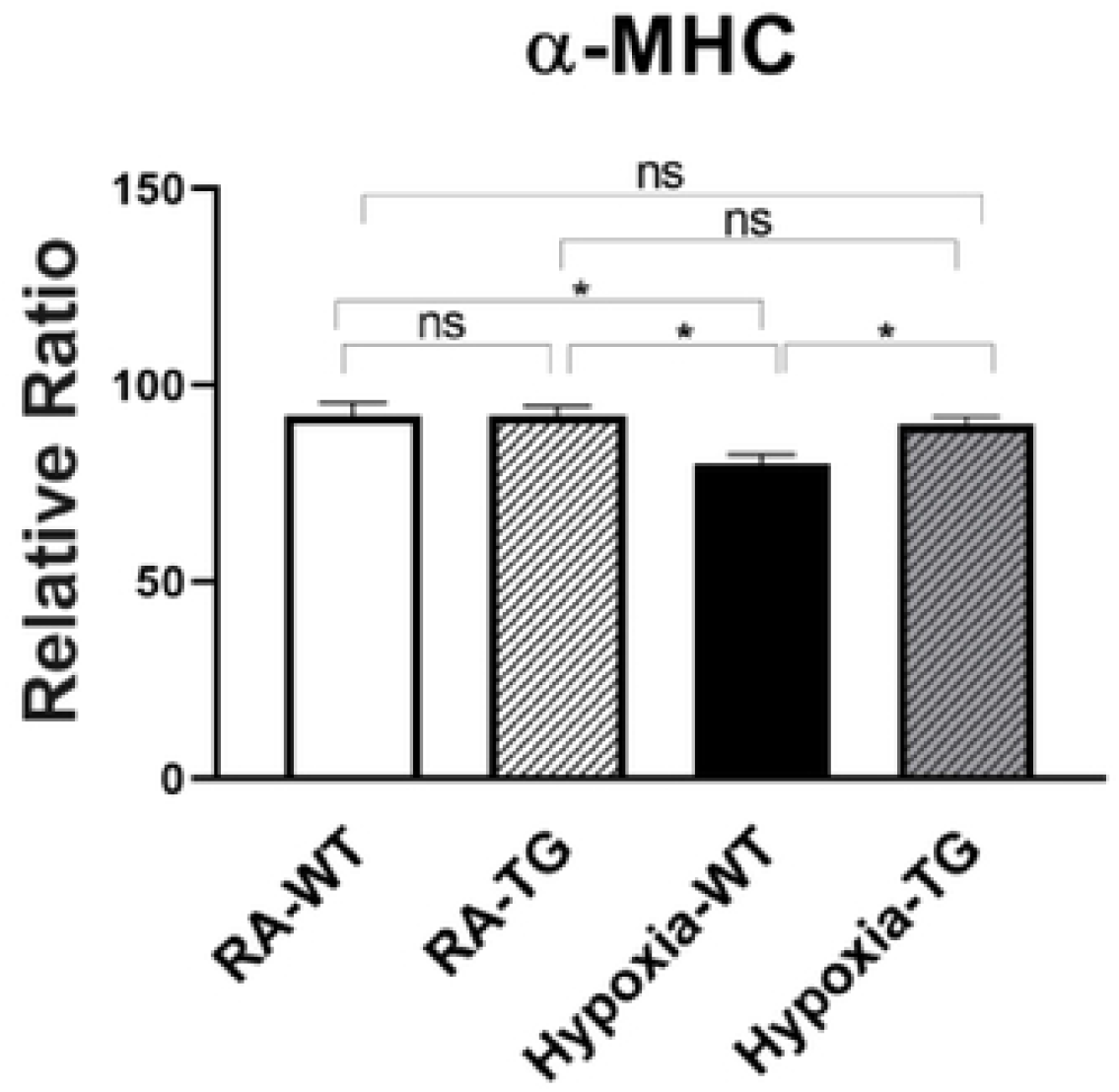
RT-PCR for α-Myosin Heavy Chain. Adult mice (C57B6) (WT) and EC-SOD transgenic mice (TG), were exposed to 10% hypoxia for 21 days (WT). Room air animals were used as control group (RA). Quantitative RT-PCR was used to assess gene expression analysis of α-MHC in the right ventricular tissue. All experiments n = 5. Data represents mean + SEM. *P:0<05, ** P< 01, *** P< 0.001, **** P < 0.0001.

#### In-vitro studies evaluating RASSF1A

Post transfection of mouse cardiac firbroblast (MCF), tissue culture incubated in room air, with SiRNA blocking the expression of RASSF1A and incubated in room air condition, both BrdU assay and cell count studies were performed. BrDU analysis (Fig. 8A) showed a significant increase in cell proliferation post-transfection with SiRNA, inhibiting RASSF1A expression, as compared to control cells incubated in room air. Cell counts (Fig. 8B) showed that compared to baseline, there was a significant increase in cell numbers, post transfection with SiRNA encoding RASSF1A, as compared to control cells transfected with an empty vector.

**Figure 8:**
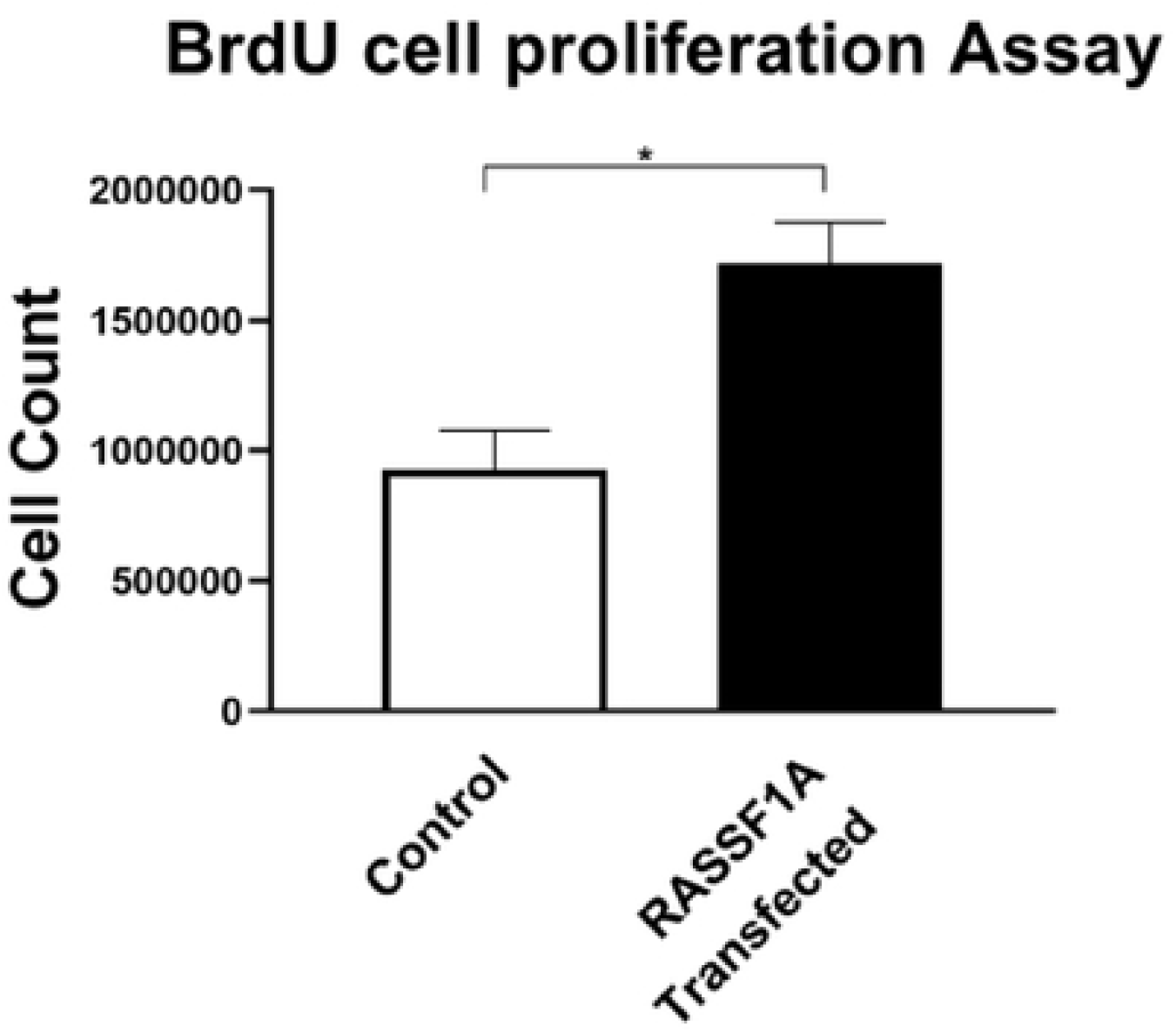
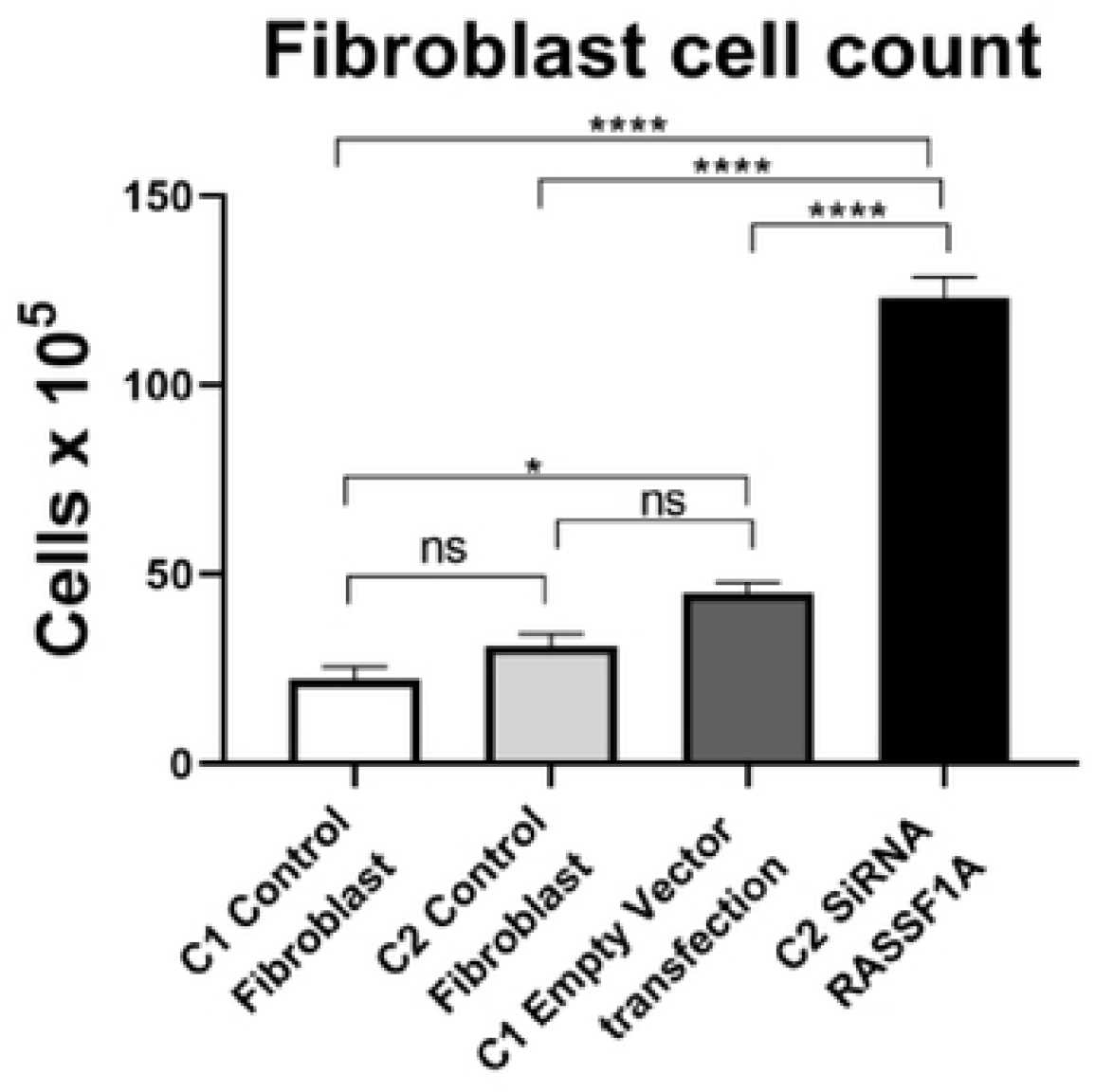
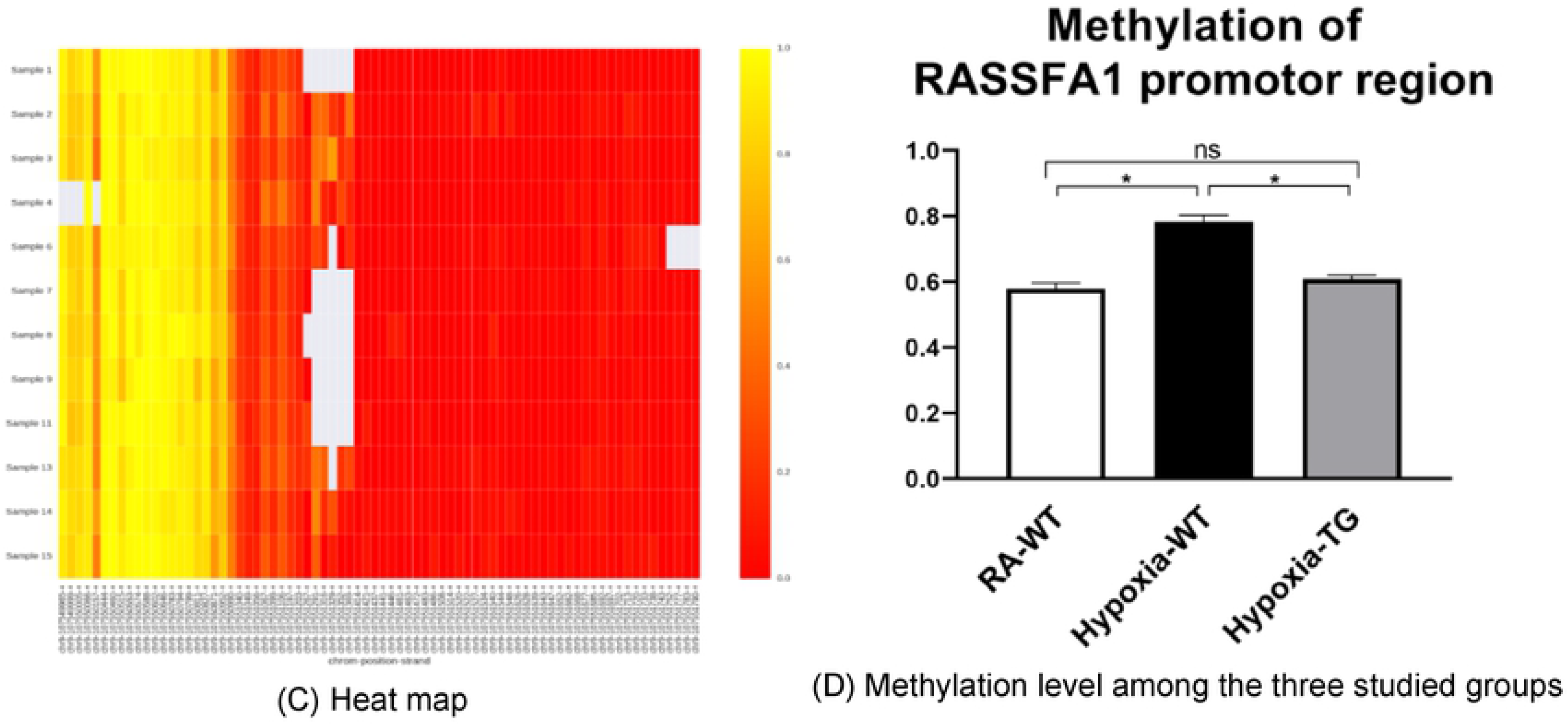
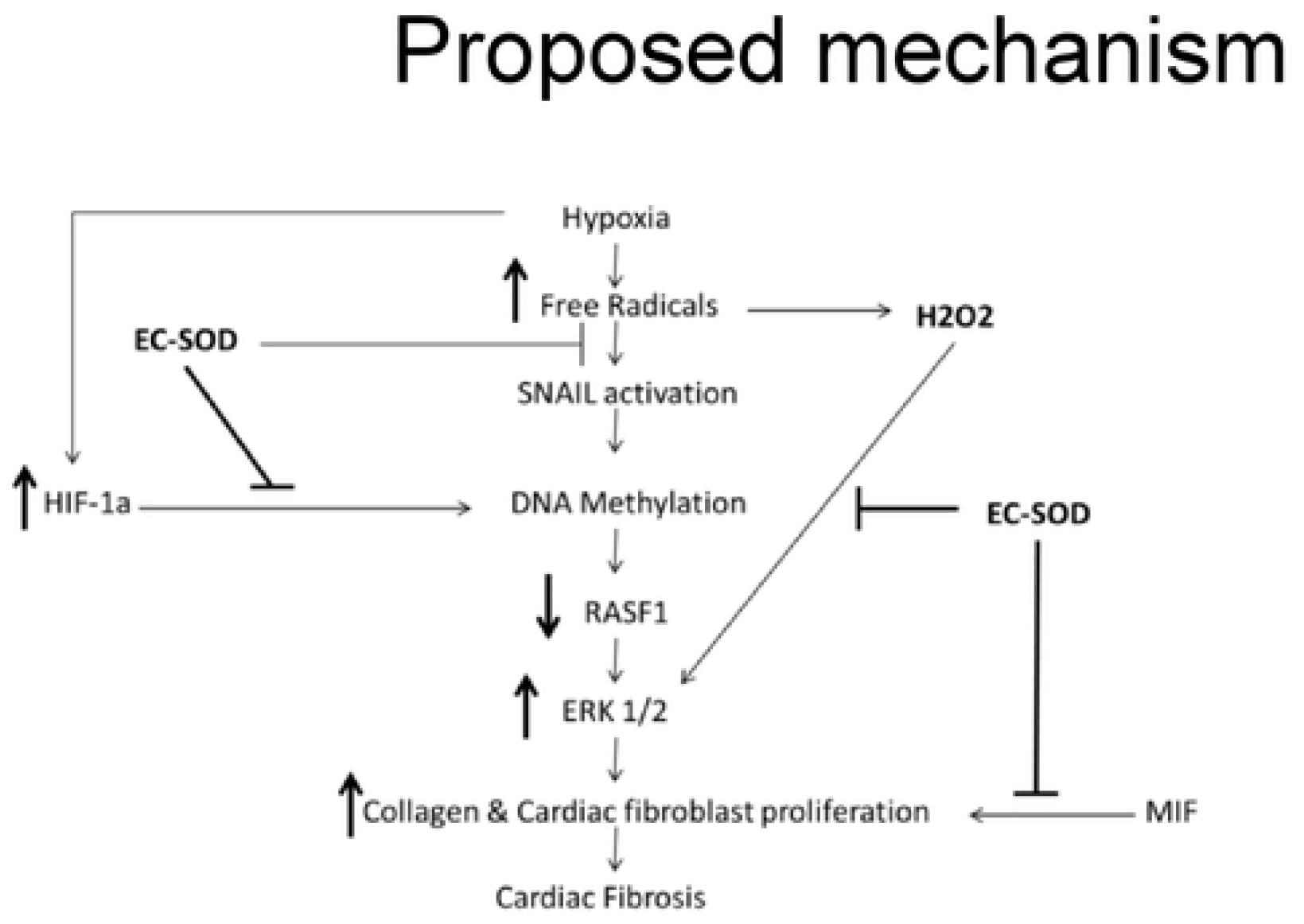
Cell Proliferation studies after transfection with SiRNA for RASSF1A –. Cardiac fibroblasts were transfected with SiRNA for RASSF1A and compared to cells placed in RA which served as a control. BrdU cell proliferation studies were undertaken (A), and cells as underwent cell counts (B). DNA methylation studies of RASSF1A promotor region as shown in the heat map (C). Qunatitative estimation of methylation level among the three studied groups (D). Proposed mechanism (E). Data represent mean + SEM. *P:0<05, ** P< 01, *** P< 0.001, **** P < 0.0001.

#### Methylation studies

mRNA expression levels of DNA methyltransferases and methyl-CpG–binding domain proteins (MBD) were studied to investigate the possible mechanism for the observed methylation differences. Evaluated by RT-PCR normalized to GAPDH, DNMT expression was significantly higher in hypoxic WT cardiac cells than the hypoxic TG cardiac cells (Figure 5A&B).

The methylation level of the RASSFA1 promotor region (8 amplicons) was significantly higher in hypoxic WT group versus hypoxic TG group (Fig. 8C&D). This suggests the methylation pattern of the RASSFA1 promoter region may be attributed to the differences in expression of methyltransferases and/or methyl binding proteins at the mRNA level.

## Discussion

Free radicals play a key role in the pathogenesis of cardiac fibrosis.^7^ Previously it has been reported that the free radical scavenger, EC-SOD, can reduce, as well as reverse, some of the changes seen secondary to chronic hypoxic stress.^14, 19^ In our study, in animals subjected to chronic hypoxia, markers of fibrosis were increased significantly including RASSFA1 promoter methylation and SNAIL which is involved in fibromodulation. Each of these changes was significantly reversed when EC-SOD was overexpressed either *in-vitro or in-vivo*. There was a significant increase in collagen 1, Collagen 3 and ASMA in hypoxic WT animals as compared to animals housed in room air (P<0.05) (Fig.1A,B,C and Fig.2A,B,C). Similar data was shown previously using human cardiac tissue.^2^ In that study, the authors showed that the degree of hypoxia was associated with increased expression of Collagen 1 and ASMA. In our animal model, overexpression of EC-SOD offered a significant protective effect, evident by reduction in the above listed fibrotic markers. Our data supports the role of oxidative insult induced by hypoxia, in the pathogenesis of cardiac fibrosis. The dismutation of these free radicals, by overexpression of EC-SOD, leads to reversal of heart pathology as shown by the immunochemistry studies (Fig. 3A&B).

The role of DNA methylation and epigenetic changes associated with cardiac fibrosis induced by hypoxia was studied previously.^3, 20^ Both DNA methyltransferase enzymes (DNMT1 and DNMT3b), which are regulated by HIF-1α (Fig. 5C); are upregulated by chronic hypoxia (Fig. 5A&B). Their upregulation was associated with increases in all fibrotic markers examined and a significant reduction in RASSF1A protein synthesis (Fig.6). In addition, the significant increase of both DNMT enzymes expression was associated with increased DNA methylation (Fig. 6). Epigenetic changes induced by prolonged hypoxia have been shown to contribute to the pro-fibrotic nature of the ischemic environment.^2^ In normal human lung fibroblasts, there is a significant global hypermethylation detected in hypoxic fibroblasts relative to normoxic controls and is accompanied by increased expression of myofibroblast markers.^19^

SNAIL gene expression is a potential target molecule in cardiac fibrosis after ischemiareperfusion (I/R), injury and or oxidative stress insult.^21^ The SNAIL gene, is best known for its capability to trigger epithelial-to-mesenchymal transition (EMT) and endothelial-to mesenchymal transition (EndMT), that may contribute to myofibroblast formation.^22^ SNAIL is activated by free radicals and mediates the actions of endogenous TGFβ signals that induce EndMT.^23^ Injection of a selective SNAIL inhibitor, remarkably suppressed collagen deposition and cardiac fibrosis in mouse I/R injury, and significantly improved cardiac function and reduced SNAIL expression *in vivo.^24^* SNAIL can recruit multiple chromatin enzymes including LSD1, HDAC1/2, and Suv39H1. These enzymes function in a highly organized manner to generate heterochromatin and promote DNA methyltransferase-mediated DNA methylation at the promoter region.^25^ Our data showed that a significant increase of SNAIL expression in WT hypoxic group was attenuated in TG animals, which may lead to a disruption of the connection between SNAIL and these chromatin-modifying enzymes, and may represent a therapeutic target for the treatment of cardiac fibrosis.

Hypoxia induced DNA methylation has been shown to be involved in regulating the process of cardiac fibrosis.^2,25^ DNA methylation mediated silencing of the RASSF1A gene which leads to up regulation of ERK1/2 that has been shown to increase cardiac fibrosis in cancer patients under chronic hypoxic stress.^4,26^ Our data showed a significant increase of ERK1/2 in the hypoxic animal group in parallel with a significant increase of DNA methylation and a reduction of RASSF1A expression (Fig. 6A). In adult cardiomyocytes, the high level of H_2_O_2_ is associated with the activation of ERK1/2 MAPK and the stimulation of protein synthesis.^27^ Increased ERK1/2 activity leads to increased cell proliferation and collagen gene expression in activated cardiac fibroblasts.^4^ Dismutation of the free radicals by the activity of EC-SOD, leads to a global decrease of DNA methylation, increased RASSF1A protein synthesis and a significant reduction in phosphorylated ERK1/2 in the transgenic hypoxic animal group (P<0.05). This finding suggest an additional contributing mechanism to cardiac dysfunction in hypoxia, which is triggered by a change in myosin heavy chain isoform.^28^ Previously it has been shown that hypoxia leads to a change in MHC from the α to β isoform which leads to decreased cardiac contractility.^27^ Free-radicals have been shown to affect this change and scavenging these free radicals by anti-oxidants can markedly attenuate cardiac fibrosis, and improve ventricular ejection fraction and fractional shortening.^28^ In our study, there was a significant reduction in the levels of α-MHC (Fig. 7) in hypoxic WT animals as compared to the hypoxia TG group (P<0.05). Hypoxia TG animals had α-MHC levels close to RA controls. This critical histological change is crucial in preserving cardiac contractility and function.

We have shown that TG animals that have an addition copy of the EC-SOD gene show a significant reduction in DNA methylation in response to chronic hypoxic stress. While further studies are needed to completely clarify the mechanism, we speculate that this reduction in DNA methylation is through a reduction in the levels of HIF-1α, which is an inducer of DNMT and hence of the process of DNA methylation.^4^ Our data show a significant reduction in HIF-1α levels in the animals that overexpress EC-SOD (P<0.05) (Fig. 5C). This suggests a mechanism by which DNA methylation leads to cardiac fibrosis. Both these changes were mitigated in our transgenic animals, which overexpress EC-SOD.

To further explore the role of RASSF1A in cardiac fibrosis, its expression was blocked in an in-vitro model using SiRNA. The presence of the SiRNA resulted in a significant increase in fibroblast proliferation (Fig. 8A&B). Furthermore, we investigated RASSF1A promoter gene methylation in the three studied groups. Our findings showed a significant increase in RASSF1A promotor region methylation in hypoxic WT group compared to normoxic animals. This methylation process was reduced by more than 10% among the hypoxic TG animals which showed a significant reduction of both biochemical and histopathological evidence of cardiac fibrosis. In summary, our study presents a novel mechanism by which EC-SOD offers cardiac protection against fibrosis induced by chronic or prolonged hypoxia. The data identifies a critical role of EC-SOD in the control of DNA methylation. Our proposed mechanism, illustrated in Fig. 8E, suggests that EC-SOD expression will chelate the free radicals, induced by hypoxia, as a result, both HIF-1α and SNAIL gene activation will be decreased and subsequently methylation enzymes activity will decrease and RASSF1A gene expression will not be silenced or decreased due to lack of methylation. RASSF1A expression downregulates the activity of the ERK1/2 pathway which regulatesactivation of both cardiac fibroblast proliferation and transition of endothelial cells to myofibroblast. Another benefit from using anti-oxidants is chelating hydrogen peroxide, which will significantly decrease the activation of ERK1/2, as it is triggered directly by hydrogen peroxide concentration. Further studies of this mechanism could lead to specific inhibition of the pathway in the clinic to significantly reduce cardiac fibrosis, and dramatically improve the outcome of this devastating condition.

## Funding

All funding was provided by institutional support.

## Conflict of interest

None

## Supplement materials

**Sup #1:** Strategy for RASSF1A sequencing.

**Sup # 2:** Methylation study design

**Sup # 3: Comparison between RA WT control group and RA TG groups:**

**3A: Collagen 1:** Comparison between RA WT and RA TG groups, no significant difference (P<0.05).
**3B: ASMA:** Comparison between RA WT and RA TG groups, no significant difference (P<0.05).
**3C: SNAIL1:** Comparison between RA WT and RA TG groups, no significant difference (P<0.05).
**3D: DNMT1:** Comparison between RA WT and RA TG groups, no significant difference (P<0.05).
**3E: DNMT 3B:** Comparison between RA WT and RA TG groups, no significant difference (P<0.05).
**3F: HIF-1α:** Comparison between RA WT and RA TG groups, no significant difference (P<0.05).
**3G: RASSF1A:** Comparison between RA WT and RA TG groups, no significant difference (P<0.05).
**3H: Phosphorylated ERK ½:** Comparison between RA WT and RA TG groups, no significant difference (P<0.05).

**Sup # 4:** Western blot of MCF transfected with human EC-SOD vs transfected with empty vector.

